# Tumor cell-derived lactic acid inhibits the interaction of PD-L1 protein and PD-L1 antibody in the PD-L1/PD-1 blockade therapy-resistant tumor

**DOI:** 10.1101/2023.08.04.551990

**Authors:** Wonkyung Oh, Alyssa Min Jung Kim, Deepika Dhawan, Deborah W Knapp, Seung-Oe Lim

## Abstract

Immune checkpoint blockade therapy targeting the PD-1/PD-L1 axis has shown remarkable clinical impact in multiple cancer types. Nontheless, despite the recent success of PD-1/PD-L1 blockade therapy, such response rates in cancer patients have been limited to tumors encompassing specific tumor microenvironment characteristics. The altered metabolic activity of cancer cells shapes the anti-tumor immune response by affecting the activity of immune cells. However, it remains mostly unknown how the altered metabolic activity of cancer cells impacts their resistance to PD-1/PD-L1 blockade therapy. Here we found that tumor cell-derived lactic acid renders the immunosuppressive tumor microenvironment in the PD-1/PD-L1 blockade-resistant tumors by inhibiting the interaction between the PD-L1 protein and anti-PD-L1 antibody. Furthermore, we showed that the combination therapy of targeting PD-L1 with our PD-L1 antibody-drug conjugate (PD-L1-ADC) and reducing lactic acid with the MCT-1 inhibitor, AZD3965, can effectively treat the PD-1/PD-L1 blockade resistant tumors. The findings in this study provide a new mechanism of how lactic acid induces an immunosuppressive environment and suggest a potential combination treatment to overcome the PD-1/PD-L1 blockade therapy resistance.

## Introduction

Immune checkpoint blockade therapy targeting the PD-1/PD-L1 axis, one of the most promising cancer immunotherapies, has shown remarkable clinical impact in multiple cancer types. However, despite the recent success of PD-1/PD-L1 blockade therapy (1-4), such impact and response rates in cancer patients have been shown to be limited (20∼40%), specifically to tumors bearing certain tumor microenvironment characteristics that bolster response to therapy. Furthermore, although PD-1/PD-L1 blockade therapy induces durable responses in cancer patients, a significant proportion of initial responders eventually develop resistance. The mechanisms leading to both primary and acquired resistance to PD-1/PD-L1 blockade therapy are varied (4-8). For example, insufficient immunogenicity of the tumor, downregulation of MHCs, T cell exhaustion, failure of interferon-gamma signaling, oncogenic signaling, altered receptor tyrosine kinase signaling, and immunosuppressive tumor microenvironment were identified or suggested as mechanisms of resistance. Several combination therapies were offered to target different steps of the cancer immunity cycle to overcome resistance, such as combining PD-1/PD-L1 blockade with chemotherapy, radiotherapy, or targeted therapy (6,7). However, the previously described mechanisms have been demonstrated to be insufficient in fully accounting for resistance.

Aerobic glycolysis is a common feature of rapidly proliferating cancer cells. Unlike normal differentiated cells, most cancer cells produce large amounts of lactic acid regardless of oxygen levels. This metabolic property is often referred to as “aerobic glycolysis” (9,10), a well-known metabolic reprogramming of cancer cells to sustain cell proliferation and a hallmark of cancer (11,12). Due to the metabolic reprogramming in cancer, the concentration of nutrients can be lower in the tumor microenvironment compared to normal tissues. Furthermore, several byproducts of the cancer cells’ metabolism may accumulate and affect the functions of immune cells (13). Of those, the most prominent metabolite in the tumor microenvironment is lactic acid. Lactic acid is transported outside the cell by monocarboxylate transporters (MCTs) and creates an acidic condition in the tumor microenvironment. Intratumoral lactic acid concentrations can reach up to 40 mM (14), and the high lactic acid concentration is known to be correlated with aggressive progression and poor survival in cancer patients (15).

Until recently, lactic acid was considered to be solely a byproduct of glycolysis. However, it was recently shown that lactic acid functions as an important regulator of cancer development and metastasis by modulating cell-to-cell interactions between cancer, stromal, and endothelial cells (16). Lactic acid has been recognized as one of the important molecules that modify immune responses in the tumor microenvironment. For example, lactic acid promotes the production of IL-17 in CD4+ Th17 cells (17), stimulates the polarization of tumor-associated macrophages into the M2-like phenotype (18), and inhibits cytotoxic CD8+ T cells (19,20). In addition, the concept of the reverse Warburg effect has been suggested as a new modality of anti-cancer treatment by preventing the generation and transport of lactic acid through the inhibition of MCTs (9,21). Our previous studies provided a link between EGF-induced extracellular lactic acid and cancer cell immune escape through the inhibition of cytotoxic T cell activity (20). Although lactic acid has been associated with the immunosuppressive tumor microenvironment, a detailed mechanism of how lactic acid modulates anti-tumor immunity in the tumor microenvironment remains unclear. Furthermore, a link between lactic acid and immune checkpoint molecules or immune checkpoint blockade therapy is still unknown.

In the current study of a new role of lactic acid in the resistance to immune checkpoint blockade therapy, we found that lactic acid levels were increased in PD-1/PD-L1 blockade-resistant tumors, and lactic acid inhibited the PD-L1 protein and PD-L1 antibody interaction. Reduction of lactic acid in resistant tumors mediated by an MCT-1 inhibitor enhanced the therapeutic efficacy of PD-1/PD-L1 blockade therapy. Hence, normalizing the altered lactic acid levels in the tumor microenvironment can improve the therapeutic efficacy of current PD-1/PD-L1 blockade therapy.

## Results

In order to study the underlying mechanisms of PD-1/PD-L1 blockade resistance, we established multiple syngeneic mouse tumor models, including mouse breast cancer cells, EMT6 and 4T1, mouse lung cancer cells, LLC1, and mouse colon cancer cells, CT26, resistant to PD-1/PD-L1 blockade therapy after three rounds of injection/treatment/isolation (Fig. 1A). Indeed, the resistant cells (R3) showed resistance to PD-1/PD-L1 blockade antibody treatment in mice (Fig. 1B). Interestingly, in our PD-1/PD-L1 blockade resistant tumor models, we found: (1) no loss of PD-L1 protein expression in the tumor cells, (2) decreased cytotoxic T cell population, and (3) increased MDSC population that suppresses cytotoxic T cells (Fig. 1C-H). These data imply that resistant tumors have an immunosuppressive tumor microenvironment while maintaining PD-L1 expression.

**Figure 1.**
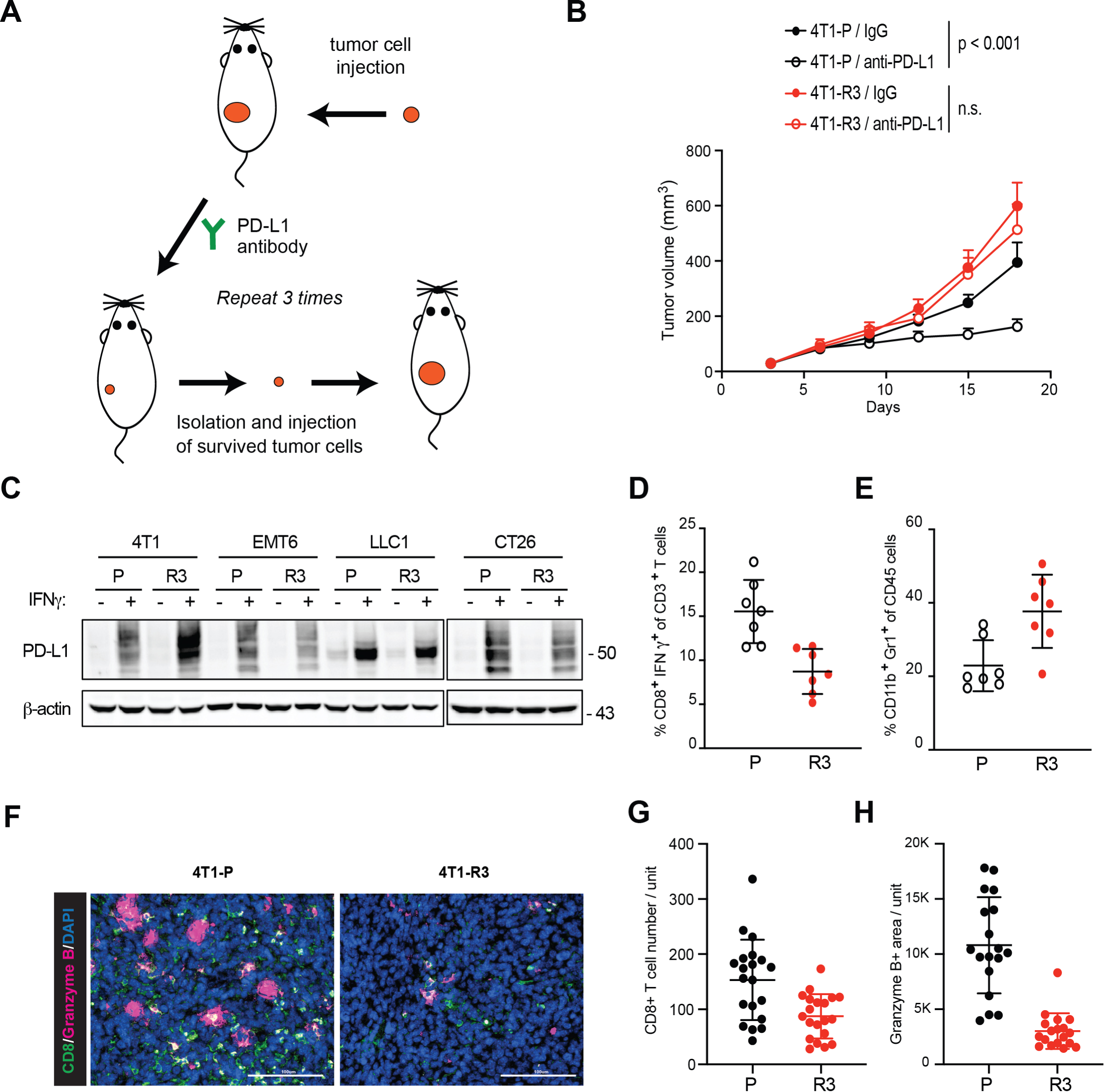
The tumor cells (R3) resistant to PD-1/PD-L1 blockade therapy. (A) Experimental strategy for establishment of PD-1 blockade therapy resistant 4T1, EMT6, E0771, LLC1, or CT26 cell lines. (B) 4T1-P (parental) and -R3 (resistant cells) tumor growth in BALB/c mice following PD-1 blockade therapy. (C) PD-L1 expression of P or R3 tumor cells were analyzed by Western blot. Tumor cells were treated with IFN gamma (IFNγ). (D) Intracellular cytokine staining of CD8^+^ IFNγ ^+^ cells in CD3^+^ T cell populations from isolated tumor-infiltrating lymphocytes. n = 8 per group. (E) MDSC (CD45^+^CD11b^+^Gr-1^+^) population was analyzed by flow cytometry. n = 8 per group. (F-H) Immunofluorescence staining of protein expression of CD8, and granzyme B in the PD-1/PD-L1 blockade resistant 4T1 tumor masses. Hoechst, nuclear counterstaining. Scale bar, 100 μm. Representative images of immunostaining of CD8 and granzyme B in the 4T1 tumor mass (F). CD8 (G) and granzyme B (H) were quantified using Gen5 software. n = 22.

In our previous study, breast cancer cells were shown to produce high amounts of lactic acid and inhibit anti-tumor immunity (20). Therefore, we asked whether lactic acid levels increased in the resistant tumors. Interestingly, the lactic acid levels were higher in the resistant tumors than that of parental tumors (Fig. 2A). Several studies, including our previous study, showed that lactic acid can suppress anti-tumor immunity by inhibiting cytotoxic T cell activity in the tumor microenvironment (20,22-24). Thus, we can deduce that high concentrations of lactic acid led to an acidic condition in the tumor microenvironment. However, the mechanism behind the immunosuppressive tumor microenvironment brought upon by the acidic condition produced by the lactic acid is not yet fully understood. Besides its suppressive effects on T cells, we have made the novel observation that lactic acid can inhibit the binding of PD-L1 antibodies to the PD-L1 protein. Specifically, lactic acid decreased the interaction between the PD-L1 protein and anti-PD-L1 antibody (Fig. 2B). The concentration of lactic acid in the resistant tumors was 3 ∼ 9 mM (Fig. 2A), and 5 ∼ 10 mM of lactic acid significantly inhibited the PD-L1 protein and PD-L1 antibody interaction *in vitro* (Fig. 2B). Therefore, the increase of lactic acid in the resistant tumors may inhibit the interaction of the PD-L1 protein with the PD-L1 antibody. These data suggest that tumor cell-derived lactic acid in the resistant tumors plays an important role in generating the immunosuppressive tumor microenvironment.

**Figure 2.**
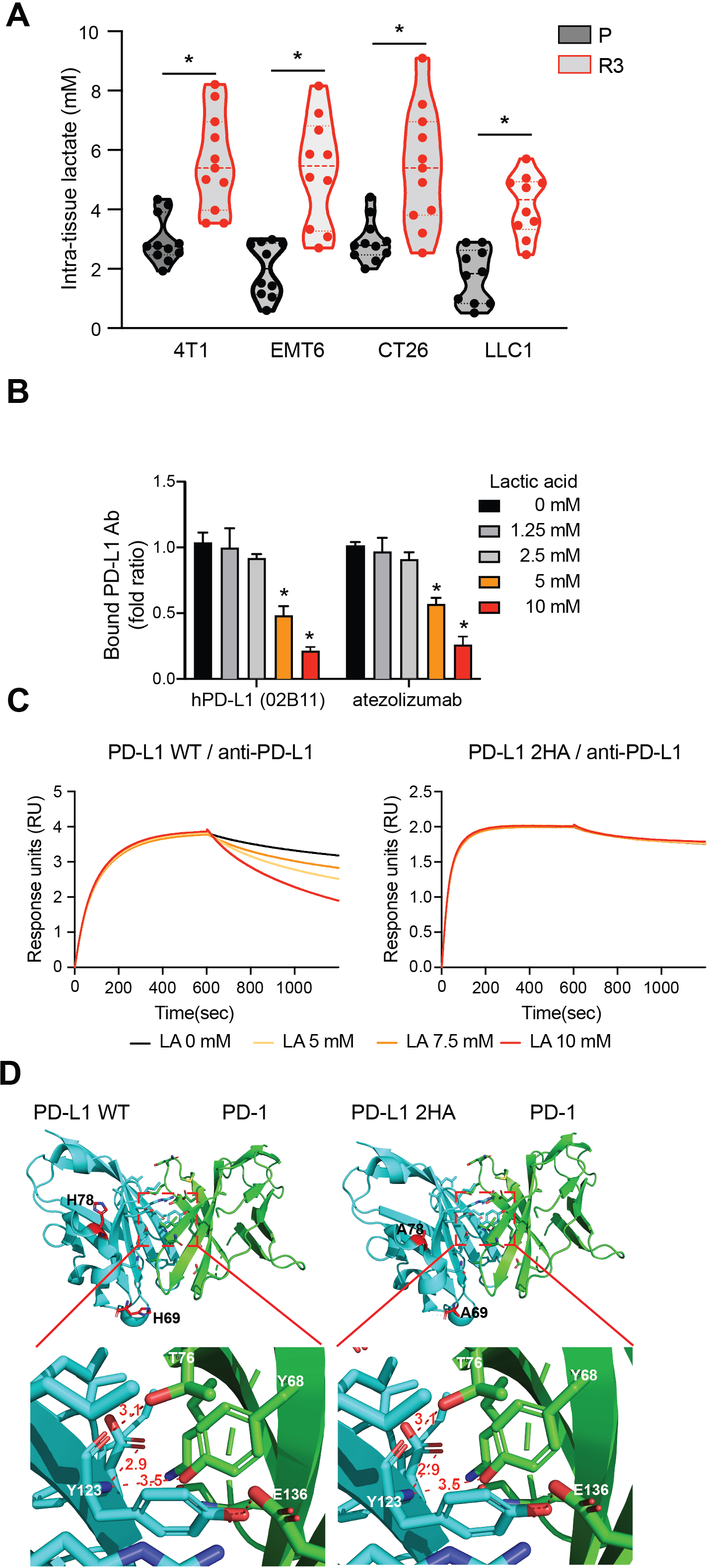
Lactic acid decreases the PD-L1 and PD-L1 antibody interaction. (A) Lactic acid increases in the resistant tumors. Intratumoral lactic acid levels in the PD-1/PD-L1 blockade resistant 4T1, EMT6, CT26, and LLC1 tumors. (B) PD-L1 protein and PD-L1 antibody interaction was determined by ELISA. (C) PD-L1 wild type (WT) or PD-L1 2HA (H69A/H78A) mutant and PD-L1 antibody binding affinity was determined by Octet. (D) The PD-1/PD-L1 WT and PD-1/PD-L1 2HA interface. The numbers represent the distance (Å) between amino acids on the PD-1/PD-L1 proteins.

The histidine residue plays a critical role in regulating the binding affinity of protein-protein interactions (or protein-antibody). Six histidine residues (H69, H78, H151, H172, H220, and H233) exist on the extracellular domain (ECD) of the PD-L1 protein. Therefore, we asked whether the lactic acid-induced acidic condition inhibits the PD-L1 protein/PD-L1 antibody interaction through the modulation of the histidine residue(s) on the PD-L1 protein. Indeed, the PD-L1 2HA (H69A and H78A) mutation abolished the lactic acid-induced decrease in PD-L1/PD-L1 antibody interaction (Fig. 2C). However, there was no change in the distance between PD-L1 and PD-1 caused by the H69A and H78A mutations (Fig. 2D). These data imply that the two histidine residues (H69 and H78) play an important role in the lactic acid-induced modulation of the PD-L1/PD-L1 antibody interaction.

Normalizing the level of lactic acid with glycolysis inhibitors can improve the therapeutic efficacy of the PD-L1 antibody in the resistant tumor models. Indeed, among several glycolysis inhibitors, an MCT-1 inhibitor, AZD3965, significantly inhibited the tumor growth of the resistant tumor cells in mice without toxicity issues (Fig. S1A). Furthermore, the level of intratumoral lactic acid was decreased (Fig. 3A) and the mRNA expression of several cytokines/chemokines (i.e., IL16, IL24, and CXCL5) that are known to enhance anti-tumor immunity was increased in the AZD3965-treated resistant tumors (Fig S1B). Given that resistant tumors do not lose PD-L1 expression and produce a high level of lactic acid in the tumor microenvironment, we hypothesized that a combination of a PD-L1 antibody-drug conjugate (PD-L1-ADC) with an MCT-1 inhibitor effectively eradicates PD-1/PD-L1 blockade therapy-resistant tumor cells. To test the hypothesis, we treated the PD-1/PD-L1 blockade-resistant tumors with the combination of PD-L1-ADC with AZD3965. The addition of AZD3965 was shown to enhance the therapeutic efficacy of the mouse PD-L1-ADC in the resistant tumor (Fig. 3B-F).

**Figure 3.**
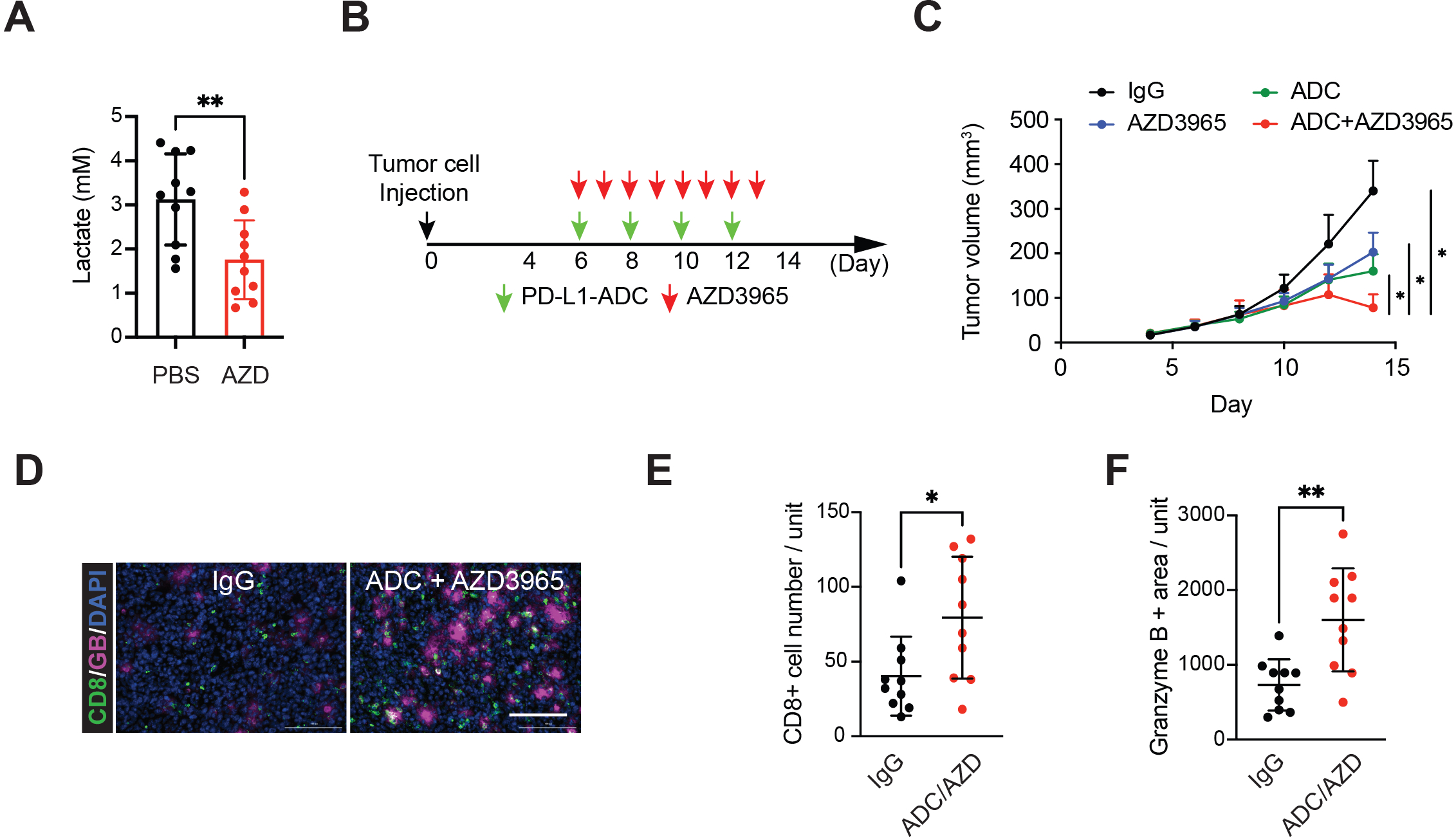
Mouse PD-L1 antibody-drug conjugate enhances anti-tumor immunity in the resistant EMT6 tumors. (A) Intratumoral lactic acid levels in the PD-1/PD-L1 blockade resistant tumors. AZD, AZD3965. (B) Schedule of drug treatments. (C) Tumor growth of resistant EMT6 tumors in BALB/c mice treated with mouse PD-L1 (MIH6)-ADC and/or AZD3965. n = 7. (D) Representative images of immunostaining of CD8 (green) and granzyme B (GB; magenta) in the IgG or PD-L1-ADC/AZD3965 treated EMT6 tumor masses. (E and F) Immunofluorescence staining of the protein expression pattern of CD8 and granzyme B in the tumor masses from IgG or PD-L1-ADC/AZD3965-treated mice.

Furthermore, we also validated the therapeutic efficacy of the combination treatment of human PD-L1-ADC with AZD3965 in the humanized PD-L1 mice. To do so, we have developed our own hPD-L1 antibodies that can be internalized and also validated their binding affinity and specificity for use in *in vivo* mouse studies (Fig. 4A). The hPD-L1 antibody (02B11 clone) recognized the human PD-L1 protein (Fig. 4A, top, and 4B) and blocked the human PD-1/PD-L1 interaction (Fig. 4A, middle). Moreover, as aforementioned, unlike other FDA-approved antibodies, atezolizumab and durvalumab, our hPD-L1 antibody can be internalized, which can be utilized in the development of antibody-drug conjugates (Fig. 4A, bottom, and 4B). To evaluate the therapeutic efficacy of our human PD-L1 antibody *in vivo*, we established humanized PD-L1 mice in which the mouse *cd274* (PD-L1) has been replaced with human PD-L1 using CRISPR knock-in mouse technologies (Fig. 4D and 4E). Also, human PD-L1 expressing E0771 cells (E0771^hPD-L1^) were established for use as a syngeneic mouse model (Fig. 4F). To enhance the cytotoxicity of the PD-L1-ADC and eradicate the tumor cells directly, MMAE was conjugated to our human PD-L1 antibody through a cleavable valine-citrulline (vc) linker (25). Similar to the E0771 tumor model (Fig 3), the combination treatment of hPD-L1-ADC with AZD3965 inhibited the tumor growth of the resistant E0771^hPD-L1^ cells and enhanced anti-tumor immunity (Fig. 4E-G). These data suggest that a combination of PD-L1-ADC with an MCT-1 inhibitor, AZD3965, effectively eradicates PD-1/PD-L1 blockade therapy-resistant tumor cells.

**Figure 4.**
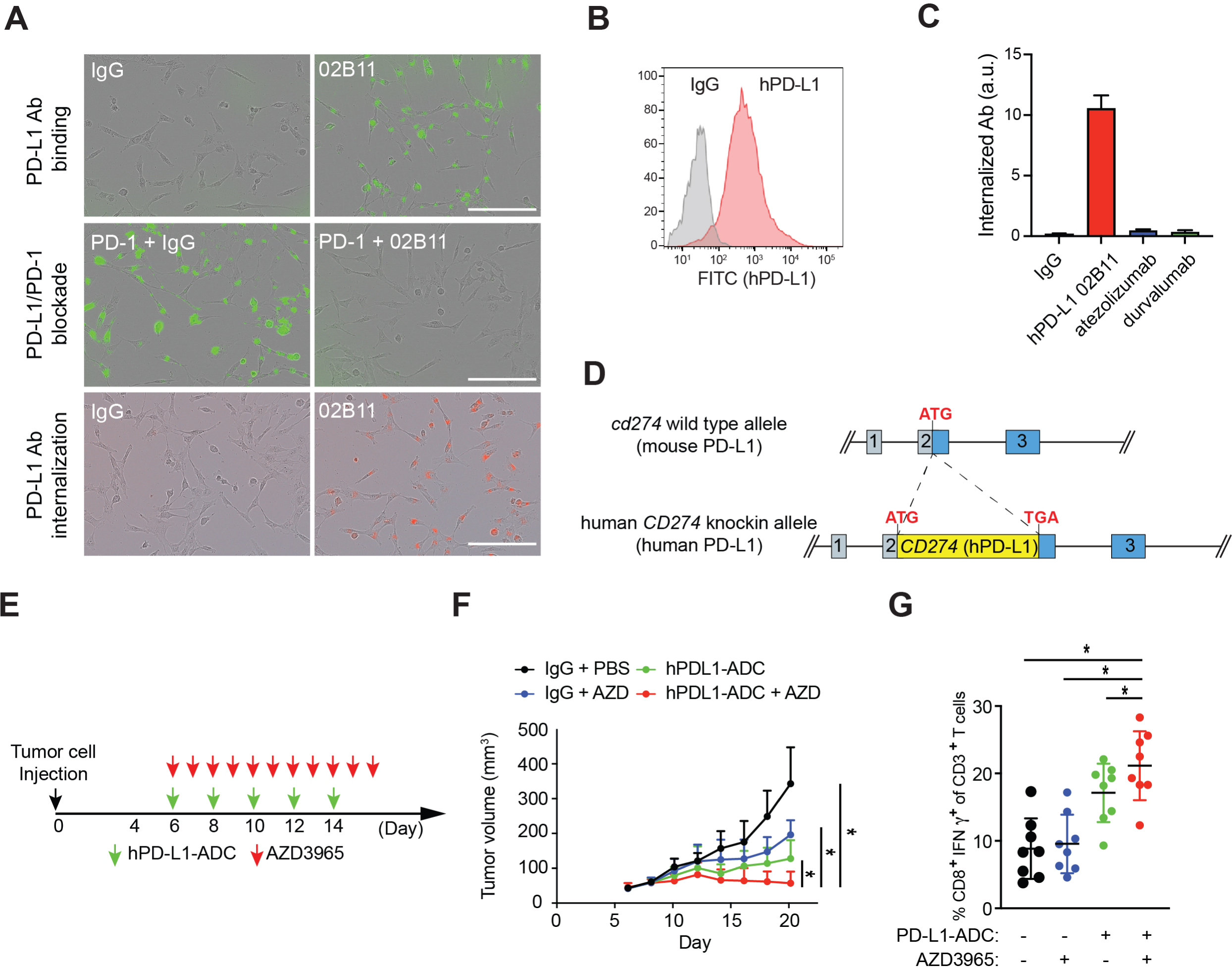
The combination of human PD-L1 02B11 antibody-drug conjugates and AZD3965 eradicates PD-L1 blockade resistant tumor cells in the humanized PD-L1 mice. (A) Representative images of the PD-L1 antibody binding (top), PD-1/PD-L1 blockade by PD-L1 antibodies (middle), and PD-L1 antibody internalization (bottom). Green fluorescence merged images of PD-L1 expressing cells are shown. 01C11 and 02B11 are clone numbers. Red fluorescence (pHrodo^TM^ red conjugated PD-L1 antibodies) represents the internalized PD-L1 antibodies. (B) Flow cytometric analysis of membrane located human PD-L1 protein on E0771^hPD-L1^ cells. (C) Internalization of hPD-L1 02B11, atezolizumab, and durvalumab. The internalized antibodies were quantified by red fluorescence. (D) Knock-in strategy for humanized PD-L1 mice. (E) Schedule of drug treatments. (F) The tumor growth of hPD-L1 expressing E0771 cells, E0771^hPD-L1^ in the humanized PD-L1 mice. *n* = 6 mice per group. (G) Intracellular cytokine staining of CD8^+^ IFNγ ^+^ cells in CD3^+^ T cell populations from isolated tumor-infiltrating lymphocytes. n = 8 per group.

## Discussion

Our current study demonstrates the effect of lactic acid in modifying immune checkpoint molecules, specifically in modifying the PD-L1 protein and anti-PD-L1 antibody interaction, in the tumor microenvironment. As a rapidly evolving field, immunotherapy targeting an immune checkpoint receptor/ligand has changed the paradigm of cancer treatment. The altered profiles of metabolites and acidification in the tumor microenvironment are well-known characteristics of immunosuppression. However, there is a gap in knowledge of the underlying regulatory mechanism of how tumor cell-derived metabolites render the immunosuppressive tumor microenvironments. We filled the gap by identifying a new role of lactic acid in the inhibition of interactions between immune checkpoint molecules and their antibodies.

The altered metabolic activity of cancer cells shapes the anti-tumor immune response by affecting the activity of immune cells. In particular, glycolytic metabolites, such as glucose and lactic acid, regulate T cell proliferation and function. However, it remains mostly unknown how the altered metabolic activity of cancer cells impacts the therapeutic efficacy of and resistance to the PD-1/PD-L1 blockade therapy. Among the altered metabolites, we found a new role of lactic acid in the tumor microenvironment that impacts the efficacy of immunotherapeutic antibodies. It is known that the binding affinity between peptides and MHC molecules changes in a pH-dependent manner via conformational changes (26). For example, low-affinity peptides strongly bound at pH 7.0 can be released at pH 5.0. The imidazole group is a critical element for such pH sensitive molecular switches, and the protonated and the nonprotonated forms of imidazole are chemically very different. The imidazole group of histidine is known to be the only amino acid side chain affected by the changes in pH (26). Thus, the histidine residue may play a critical role in regulating the binding affinity of protein-protein interactions (or protein-antibody) in the tumor microenvironment. Indeed, the lactic acid-induced decrease in PD-L1 protein/PD-L1 antibody interaction relied on the two histidine residues, H69 and H78, on the extracellular domain of the PD-L1 protein (Fig. 2). These data imply that the acidification of the tumor microenvironment may alter the interactions between immune receptors with their ligands or the therapeutic antibodies that targeting them via the modulation of the histidine residues on the immune receptors.

Targeting PD-L1 with an ADC in breast cancer is well justified (27), as the protein expression of PD-L1 in the resistant tumor cells was validated in multiple cancer types (i.e. breast, lung, and colon cancers; Fig. 1C). The PD-1/PD-L1 therapy resistant syngeneic mouse tumor cells and the humanized PD-L1 mice are appropriate preclinical models to test the therapeutic efficacy of our human PD-L1-ADC and other potential drugs in. In the development of immuno-oncology drugs, particularly immunotherapeutic antibodies, translation of the discoveries from mouse models to clinical trials has been hindered by many biological differences between mice and humans, such as the lack of cross-reactivity between species. The lack of cross-reactivity is one of the major obstacles for developing human immune checkpoint blockade antibodies to further translate those with the highest success in mice into human trials. To overcome this obstacle, we established the humanized PD-L1 mouse model as a preclinical tool and utilized it in validating the therapeutic efficacy of immunotherapeutic PD-L1 antibodies including our own PD-L1 antibody and other FDA-approved antibodies (i.e. atezolizumab, durvalumab). Our humanized PD-L1 mouse model and syngeneic mouse breast cancer cell line, E0771 ^hPD-L1^, is a unique and powerful tool for preclinical immuno-oncology research.

Collectively, we uncovered a new mechanism of how tumor cells create an immunosuppressive microenvironment by altering lactic acid production. Our findings also suggest a new combination treatment to improve the efficacy of current immune checkpoint blockade therapies. Our studies, therefore, provide the preclinical data necessary for the development of new treatment strategies capable of increasing cancer survival rates by enhancing the therapeutic efficacy of immune checkpoint blockade therapies.

## Materials and Methods

### Cell culture, stable transfectants and transfection

4T1, EMT6, E0771, LLC1, and CT26 mouse cancer cell lines and BT549 human breast cancer cell lines were obtained from ATCC (Manassas, VA, USA) and Millipore Sigma (St. Louis, MO, USA), respectively. Cells were grown in DMEM or DMEM/F12 medium supplemented with 10% fetal bovine serum. Using a pGIPZ-shPD-L1/Flag-hPD-L1 dual-expression construct to knockdown endogenous mouse PD-L1 and reconstitute Flag-hPD-L1 simultaneously (28), we established endogenous PD-L1 knockdown and Flag-hPD-L1 expressing E0771 cell lines. Lentivirus was packaged by co-transfecting transfer plasmids with pMD2.G (Addgene #12259) and pCMV dR8.2 (Addgene #12263) to Lenti-X^TM^ 293 cells (Takara Bio, San Jose, CA, USA) with X-tremeGENE HP (Roche Diagnostics, Indianapolis, IN, USA), and the supernatant was harvested for lentiviral transduction. Selection with 1 μg/mL puromycin (InvivoGen, San Diego, CA, USA) was routinely performed to maintain ectopic gene expression. For mouse PD-L1 knockout, we transfected mouse PD-L1 double nickase plasmid (Santa Cruz Biotechnology, Dallas, TX, USA) into E0771 cells using X-tremeGENE transfection reagent. For human PD-L1 overexpression E0771 cells (E0771^hPDL1^), we infected mouse PD-L1 KO E0771 cells with lentivirus carrying pGIPZ-Flag-hPD-L1 followed by selection with puromycin.

### Creation and selection of anti-human PD-L1 monoclonal antibodies

Anti-human PD-L1 monoclonal antibody, 02B11 was generated via conventional hybridoma procedures using BALB/c mice immunized with the extracellular domain of hPD-L1 (attached to a 6xHis tag; Novoprotein Scientific Inc., Beijing, China) at Purdue University. Splenocytes were isolated from the immunized mice and then fused with SP2/0 myeloma cells. Supernatants from isolated clones were screened for the ability to block the PD-1/PD-L1 interaction through human PD-L1 expressing cell-based assays. Clonal antibodies were purified from supernatants and the same assays were rerun.

### Generation of the human *CD274* knock-in mouse

The humanized PD-L1 mouse (human *CD274* knock-in mouse) was generated by *Easi*-CRISPR (Efficient additions with ssDNA insert-CRISPR) strategy using a long single-strand DNA (ssDNA) donor and CRISPR ribonucleoproteins (29). Briefly, the long ssDNA (a full length of human *CD274* cDNA; NM_014143.4) was injected with pre-assembled guide RNA (gRNA, CAGCAAATATCCTCATGTTT TGG) and Cas9 ribonucleoprotein (ctRNP) complexes into mouse zygotes. The ssDNA and sgRNA were synthesized at Integrated DNA Technologies (IDT, Coralville, IA, USA). All animal experiments for the knock-in mouse generation performed were approved by the Purdue Animal Care and Use Committee (PACUC) at Purdue University. C57BL/6N female mice at 4 weeks of age (Envigo, Indianapolis, IN, USA) were superovulated and then mouse zygotes were obtained by mating C57BL/6N males with the superovulated females. Pronuclei of one-cell stage fertilized mouse embryos were injected with 20 ng/μl Cas9 protein, 10 ng/μl sgRNA, and 5 ng/μl ssDNA. Microinjections and mouse transgenesis were performed as described (30). Mouse genomic DNA was extracted from the tail tip and then used for the genotyping (Primer set 1 forward, 5’-CCACTTGGTTCTACATGGCT -3’; Primer set 1 reverse, 5’-GTGACTGGATCCACAACCAA -3’; Primer set 2 forward, 5’-CCATCAAGTCCTGAGTGGTAAG -3’; Primer set 2 reverse, 5’-GGACTAAGCTCTAGGTTGTCC-3’; Primer set 3 forward, 5’-GACTGGCTTTTAGGGCTTATGT -3’; Primer set 3 reverse, 5’-ACACCCCACAAATTACTTCCATT -3’) and sequencing (Primer set 3 forward, 5’-GACTGGCTTTTAGGGCTTATGT -3’; Primer set 3 reverse, 5’-ACACCCCACAAATTACTTCCATT -3’) to verify the location of insertion and DNA sequence of human *CD274*.

### Mouse study and antibody treatment

All procedures with BALB/c, C57BL/6, or the humanized PD-L1 mice (C57BL/6 strain; 6- to 8- week-old) were conducted under guidelines approved by the PACUC at Purdue University. Mice were divided according to the mean tumor volume in each group. 4T1, EMT6, LLC1, CT26, or E0771 ^hPD-L1^ (2 × 10^5^ cells in 25 μL of medium mixed with 25 μL of Matrigel Basement Membrane Matrix [BD Biosciences, San Jose, CA, USA]) were injected into the mammary fat pad or flank. For treatment with antibodies, 5 mg/kg of hPD-L1 antibody (02B11 clone or atezolizumab-mIgG2a), mouse PD-L1 (10F.9G2 clone [BioXcell, Lebanon, NH, USA] or MIH6 clone [BioLegend, San Diego, CA, USA]), control mouse IgG, or control rat IgG (Bio X Cell) was injected intraperitoneally on days 6, 8, 10, 12, and 14 after tumor cell inoculation when tumor size was approximately 30∼40 mm^3^. Tumors were measured every other day with a caliper, and tumor volume was calculated using the following formula: π/6 × length × width^2^.

To establish the cells resistant to PD-1/PD-L1 blockade therapy, we injected 100,000 mouse tumor cells per mouse into the mammary fat pad or flank. After 5 to 6 days, we treated the mice with PD-L1 therapeutic antibodies (7.5 mg/kg; mouse PD-L1 10F.9G2 clone or atezolizumab-mIgG2a), intraperitoneally every other day for two weeks. On day 18, we isolated the tumor cells. To enrich the resistant population, we repeated two rounds of implantation and PD-L1 antibody treatment.

### Immunofluorescence study of mouse tumor tissues

Tumor masses were frozen in an OCT block immediately after excision. Cryostat sections of 5-µm thickness were attached to saline-coated slides. Cryostat sections were fixed with 4% paraformaldehyde for 30 minutes at room temperature and blocked with blocking solution (1% bovine serum albumin, 2% donkey and/or chicken serum, and 0.1M PBS) at room temperature for 30 minutes. Samples were stained with primary antibodies against CD8 and granzyme B overnight at 4D, followed by secondary antibodies at room temperature for 1 hour. Nuclear staining was performed with Hoechst 33342 (ThermoFisher Scientific, Wlathan, MA, USA). The stained sections were visualized by automated microscopy (Lionheart LX; BioTek Instruments, Inc., Winooski, VT, USA). Granzyme B positive area and the number of CD8 positive CTL were assessed per high power field (200X). Fourteen randomly chosen microscope fields from 4 serial sections in each tissue block were examined for the number of CD8 positive CTL and granzyme B positive areas for each tissue.

### Nanostring analysis

RNA was isolated from tumor mass previously (RNeasy kit, Qiagen, Germantown, MD) and submitted to IU Research Core (IU Research Core, IUPUI, Indianapolis) for detection of modulation of genes upon activation using nCounter® mouse PanCancer Immune Profiling Panel (Nanostring Technologies, Seattle, WA). Data were analyzed using Rosalind (Rosalind, San Diego, CA). Groupwise comparison was conducted using control IgG treated tumors (n = 3) and compared with AZD3965 treated tumors (n=3). Differentially expressed genes (FC≥ 1.5; p<0.05) were considered significant. Data were visualized using heatmap, volcano plot and histogram for specific genes.

### Expression and purification of a recombinant human PD-L1 antibodies

The codon-optimized for CHO variable light and heavy chains (02B11, atezolizumab, and durvalumab) were synthesized (ThermoFisher Scientific) and then cloned into pTRIOZ-mIgG2a/mkappa (κ) vector (InvivoGen, San Diego, CA, USA). Plasmids encoding hPD-L1 02B11, atezolizumab, durvalumab antibodies: pTRIOZ-mIgG2a/mκ-02B11, pTRIOZ-mIgG2a/mκ-atezolizumab, pTRIOZ-mIgG2a/mκ-durvalumab, respectively, were transfected into ExpiCHO-S cells following the transfection kit instructions (GIBCO, A29133). ExpiCHO-S cells were cultured with ExpiCHO Expression Medium (ThermoFisher Scientific) in a shaker incubator set at 120 rpm, 37 ::JC and 8.0% CO2. Cells were collected 10 days post-transfection at 4,000 x g and 4 °C for 20 min. The antibody supernatant passed through a 0.22-µm filter and neutralized with 10XPBS buffer (Lonza™ BioWhittaker™ Phosphate Buffered Saline (10X), BW17-517Q). The antibody supernatant were pre-incubated with protein G agarose for 2 hrs. The agarose A-conjugated antibody were applied to the column (BioRad poly-prep chromatography column, #731-1550). The column was washed with low-endotoxin PBS (Lonza™ BioWhittaker™ Dulbecco’s Phosphate Buffered Saline (1X) w/o Calcium and Magnesium, BW17512F24). The bound antibody were eluted with elution buffer (ThermoFisher Scientific, Elution Buffers, 0.1M Glycin-HCl, pH2.8, #21004) into Neutralization Buffer (Tris HCl, 1M, BP1757-500). The purified antibody was concentrated and buffer exchanged with PBS, pH7.0. The antibody concentration was determined by UV absorbance at 280nm.

### The cell free PD-L1 protein/PD-L1 antibody binding and PD-L1/PD-1 blockade assays

Enzyme-linked immunoassay (ELISA) based assays were performed to compare the receptor/ligand and receptor/antibody binding. The 6X His tagged extracellular domain of hPD-L1 WT or 2HA proteins were expressed in the ExpiCHO cell system (ThermoFisher Scientific) and purified by the Ni-NTA agarose (ThermoFisher Scientific) according to the manufacturer’s protocol. For the PD-L1 protein/PD-L1 antibody binding assay, Pierce Ni-NTA coated 96-well plates (ThermoFisher Scientific) was coated with hPD-L1-His protein and anti-PD-L1 antibody and anti-mouse IgG specific HRP conjugated secondary antibodies (SouthernBiotech, Birmingham, AL, USA) were added. The bound PD-L1 antibody was quantified by measuring OD_450_ vale with a Synergy LX multi-mode reader. For the PD-L1/PD-1 blockade assays, Pierce Ni-NTA coated 96-well plates (ThermoFisher Scientific) was coated with PD-L1-His protein and PD-1-hFc protein (human Fc protein conjugated; SinoBiological US, Wayne, PA, USA) and anti-human IgG Fc specific HRP conjugated secondary antibodies (ThermoFisher Scientific) were added. And then the PD-L1 antibodies were added. The bound PD-1-Fc protein was quantified by measuring OD_450_ vale with a Synergy LX multi-mode reader.

### The cell base PD-L1/PD-L1 antibody binding and PD-L1/PD-1 blockade assays

The antibody binding and blockade assays were performed as described previously (27,31). Briefly, to measure PD-L1 protein and PD-L1 antibody interaction, we seeded 1x10^4^ BT549 ^hPD-L1^ cells per well in 96-well plates and then incubated the plates with IgG control (Rockland Immunochemicals, Pottstown, PA, USA), anti-PD-L1 antibody, and anti-mouse Alexa Fluor 488 dye conjugate (SouthernBiotech). Every hour, green fluorescent signal was measured and quantified by IncuCyte S3 (Sartorius, Goettingen, Germany). To measure PD-1 protein on the cells, we seeded 1x10^4^ BT549 ^hPD-L1^ cells per well in 96-well plates and then incubated the plates with IgG control (Rockland Immunochemicals, Pottstown, PA, USA), 02B11 antibody, PD-1-hFc protein (human Fc protein conjugated; SinoBiological US), and/or anti-human Alexa Fluor 488 dye conjugate (ThermoFisher Scientific). Every hour, green fluorescent signal was measured and quantified by IncuCyte S3 (Sartorius, Goettingen, Germany). The Image analysis was performed according to the manufacturer’s protocol.

### Flow cytometry analysis

E0771 or E0771^hPD-L1^ cells were washed twice with ice-cold cell staining buffer (BioLegend) and stained with IgG control, mouse PD-L1 (10F.9G2), human PD-L1 (02B11) for 1 hr at 4 °C. After three washes with staining buffer, cell samples were stained with Alexa Fluor 488-conjugated anti-mouse IgG specific secondary antibody for 30 min at 4 °C. Cell samples were loaded on BD LSRFortessa (BD, Franklin Lakes, NJ, USA) for analysis. Data analysis was performed on FlowJo v9 software (BD). For tumor-infiltrating lymphocyte profile analysis, excised tumors were dissociated as a single cell using the gentleMACS Dissociator (Miltenui Biotec Inc., San Diego, CA, USA) with the mouse Tumor Dissociation kit (Miltenui Biotec) and lymphocytes were enriched on a Ficoll gradient (Sigma-Aldrich). T cells were stained using anti-CD3-Alexa Fluor 488, CD4-Alexa Fluor 647, CD8a-Alexa Fluor 594, CD45.1-APC/Cy7, IFNγ-PerCP/Cy5.5, and FoxP3-Pacific Blue antibodies. All antibodies for flow cytometry analysis were purchased from BioLegend. Stained samples were analyzed using a BD LSRFortessa (BD Bioscience) cytometer.

### Binding affinity (K_D_) determination

The binding affinity (K_D_) of PD-L1 protein/PD-L1 antibody was determined by Octet Biolayer interferometry (BLI) using the Octet RED384 system (Sartorius, Bohemia, NY, USA). Briefly, His-tagged PD-L1 protein was loaded on the Octet NTA biosensor at a concentration of 100 nM. The association step was performed by submerging the sensors in three concentrations of the anti-PD-L1 antibody (50, 100, 200 nM) in the kinetic buffer. Dissociation was performed and monitored in fresh kinetic buffer. Data were analyzed with Octet Analysis HT software (Sartorius).

### Statistical analysis

All quantitative results were displayed as the mean ± SD, with at least three biological replicates. The inter-group statistical significance was calculated by two-tail Student’s t-test. *p* < 0.05 was considered statistically significant.

### Data availability statement

The data generated in this study are available upon request from the corresponding author.

### Authors’ Contributions

W. Oh: Data curation, formal analysis, validation, visualization, methodology, writing-review and editing. A.M.J. Kim: Data curation, formal analysis, visualization, writing-review and editing. D. Dhawan: Data curation, formal analysis. D.W. Knapp: Resources, data curation, validation, writing-review and editing. S.O. Lim: Conceptualization, resources, data curation, supervision, funding acquisition, validation, investigation, visualization, methodology, writing-original draft, writing-review and editing.

## Acknowledgements

This study was funded in part by American Cancer Society RSG-22-137-01-IBCD; Ralph W. and Grace M. Showalter Research Trust grant; Purdue University Center for Cancer Research Transgenic and Genome Editing Facility supported by NCI CCSG CA23168 to Purdue University Center for Cancer Research; Purdue University Center for Cancer Research NIH grant number P30 CA023168; Purdue University Institute for Drug Discovery. The authors acknowledge the use of the Chemical Genomics Facility, a core facility of the Purdue Institute for Drug Discovery and the NIH-funded Indiana Clinical and Translational Sciences Institute.

## Notes

### Competing Interest Statement

The authors have declared no competing interest.

