## Supplementary material for "Tumor cell-derived lactic acid inhibits the interaction of PD-L1 protein and PD-L1 antibody in the PD-L1/PD-1 blockade therapy-resistant tumor": Figure S1

**Supplementary information**

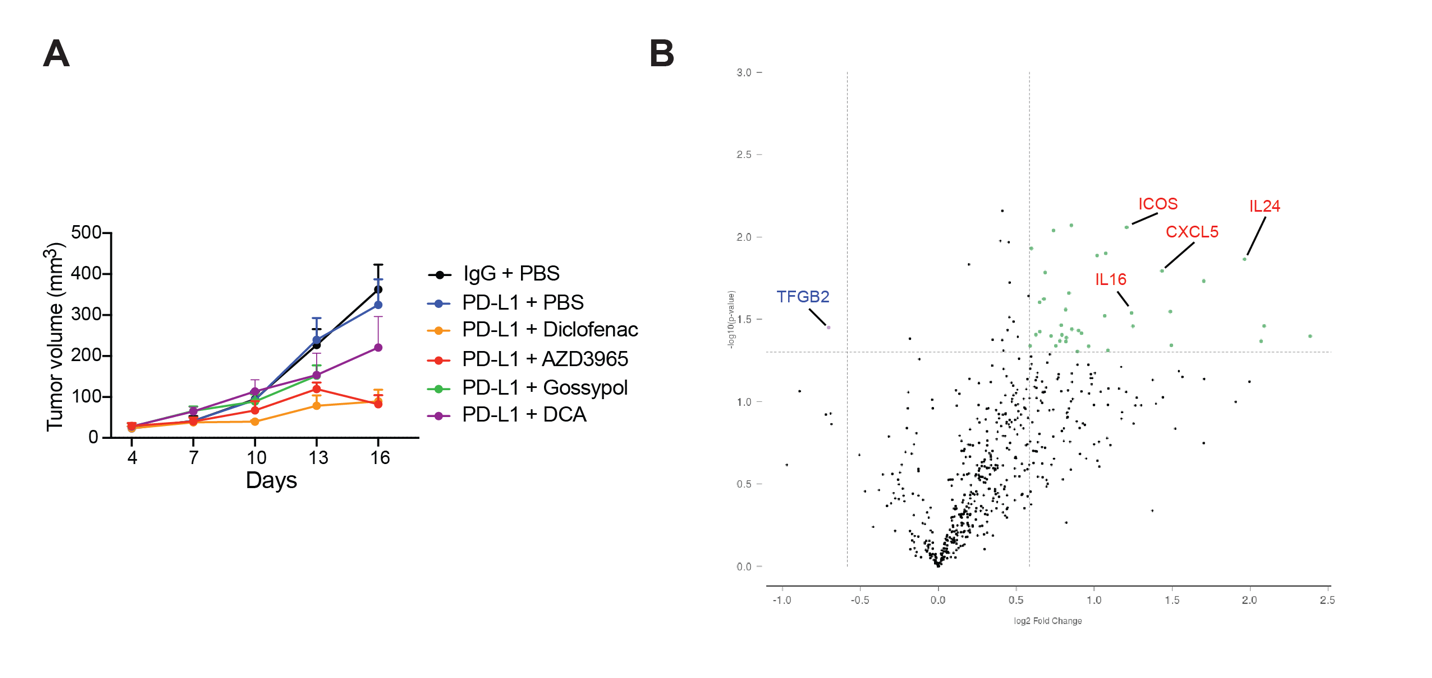

**Figure S1.** Glycolysis inhibitors inhibit a growth of PD-1/PD-L1 blockade therapy resistant tumor upon PD-1 blockade antibody treatment. (A) E0771 resistant tumor growth in the mice with PD-1 blockade therapy. aPD-L1, anti-PD-L1 antibody treatment, n = 7. (B) The differential gene expression in the AZD3965 treated tumors. The differential expression analysis was performed with Nanostring nCounter mouse PanCancer Immune Profiling Panel and Rosalind.
